# Linkage disequilibrium scaling improves robustness and power to detect genomic regions under selection

**DOI:** 10.64898/2026.01.19.700334

**Authors:** Petri Kemppainen, Frédéric Guillaume

## Abstract

Understanding the genetic basis of adaptive evolution is central to predicting how populations respond to environmental change. Genome scans and genotype–environment association methods are widely used to detect loci under selection, but because they often ignore correlations among loci (linkage disequilibrium, LD), their power decreases with increasing genetic structuring and demographic complexity. While LD contains valuable information about selection and can reduce dimensionality by summarizing signal across correlated markers, existing LD-based approaches are computationally demanding and rely on ad hoc threshold choices.

Here, we introduce a novel LD-scaled association statistic, *F* ^′^, which incorporates local LD directly into genotype–environment association tests using a fast permutation-based quantile transformation. Using forward-in-time simulations with known ground truth, we evaluate *F* ^′^ across two widely used methods—latent factor mixed models (LFMM) and EMMAX—and assess performance at the level of outlier regions rather than individual SNPs. We further integrate uncertainty in parameter choice and inference method using a consistency-based framework.

Across simulations, LD-scaling and joint inference substantially increased detection power and robustness, yielding up to an order-of-magnitude improvement under strong genetic structuring and nearly doubling performance on average across scenarios. Applying this framework to three- and nine-spined stickleback datasets, we recover well-established genomic regions associated with parallel marine–freshwater adaptation with markedly reduced background noise compared to standard analyses. Together, our results demonstrate that explicitly leveraging LD structure and integrating over analytical uncertainty provides a powerful and computationally efficient extension to genome scans for selection, improving robustness and interpretability in both simulated and empirical settings.

## 1 Introduction

Understanding the genetic basis of adaptive evolution is central to predicting how natural populations respond to environmental change (Gienapp et al., 2008; Hoffmann and Sgrò, 2011; Savolainen et al., 2013; Catullo et al., 2019). Genomic regions under selection are commonly identified using outlier ap-proaches based on population genetic statistics (Lewontin and Krakauer, 1973; Lotterhos and Whit-lock, 2015; Hoban et al., 2016; Ravinet et al., 2017; Fang et al., 2020, 2021), genotype–environment associations (Gautier, 2015; Caye et al., 2019; Forester et al., 2018; Carvalho et al., 2021), or allele fre-quency changes observed in experimental or temporal data (Long et al., 2015; Schlötterer et al., 2015; Kemppainen et al., 2017; Kelly and Hughes, 2018). Genome-wide association (GWA) and quantitative trait locus (QTL) mapping provide complementary frameworks by directly testing genotype–phenotype associations (Colosimo et al., 2005; Heerwaarden et al., 2015; Kemppainen et al., 2021).

Despite their widespread use, these approaches share common statistical limitations. In particular, they can generate false positives under complex demographic histories (Lotterhos and Whitlock, 2015; Lotterhos, 2019; Hoban et al., 2016) and often lack power when loci are treated as independent tests, leading to overly conservative multiple-testing corrections that ignore linkage disequilibrium (LD) among markers (Li et al., 2018).

Patterns of LD themselves provide valuable information about selection (Kemppainen et al., 2015; Nielsen et al., 2005; Gompert et al., 2022; Fang et al., 2020, 2021). Selective sweeps can generate tran-sient LD signatures (Smith and Haigh, 1974; Nielsen et al., 2005), while persistent local LD can arise from reduced effective recombination and gene flow between locally adapted populations, producing ge-nomic islands of differentiation (Feder and Nosil, 2010; NOSIL et al., 2009). Because population struc-ture and drift elevate LD genome-wide, however, background LD can obscure selection signals. Empir-ical and theoretical work has nevertheless shown that LD strength declines with distance from causal variants (Gompert et al., 2022), suggesting that contrasting local LD patterns against background lev-els can help focus inference on genomic regions more likely to be under selection. This perspective nat-urally motivates treating clusters of correlated SNPs as test units rather than individual markers, re-ducing dimensionality while retaining biologically meaningful signal (Kemppainen et al., 2015; Li et al., 2018; Fang et al., 2020, 2021).

Linkage disequilibrium network analysis (LDna) demonstrated that LD-based clustering can reveal complex selection signatures missed by SNP-level analyses (Fang et al., 2020, 2021). However, LDna is computationally demanding, relies on multiple user-defined thresholds, and has not been systematically evaluated using simulations with known ground truth. Here, we address these limitations by explicitly scaling test statistics with local LD patterns, combined with a permutation-based quantile transformation that controls false positives. Using genetically explicit, forward-in-time simulations implemented in Nemo (Guillaume and Rougemont, 2006), we evaluate whether this approach improves detection of loci under selection across a wide range of demographic and selective scenarios.

Because selection signals typically extend across multiple linked loci, we assess the performance at the level of *outlier regions* (ORs)—clusters of correlated SNPs reflecting shared LD with underlying causal variants—rather than individual markers, using a modified precision–recall framework. We compare standard and LD-scaled versions of two widely used association methods: latent factor mixed models (LFMM) (Frichot et al., 2013; Frichot and François, 2015; Caye et al., 2019) and the linear mixed model EMMAX (Kang et al., 2010). LFMM provides a strong benchmark for outlier detection (Lotterhos and Whitlock, 2015; Lotterhos, 2019), whereas EMMAX has been widely used in GWA studies and LD-based analyses due to its computational efficiency (Yang et al., 2014; Korte et al., 2012; Fang et al., 2021; Li et al., 2018).

Finally, because both LD-scaling and OR definition depend on multiple tuning parameters, and because different methods capture complementary aspects of genomic differentiation, we introduce a consistency-based framework that integrates evidence across parameter values and inference methods. We evaluate this approach using simulations and empirical datasets from two marine–freshwater adapted sticklebacks with contrasting population structure (Fang et al., 2021). Together, these analyses demonstrate how explicitly leveraging LD structure and integrating over uncertainty improves the robustness and interpretability of genome scans for selection.

## 2 Materials and Methods

### 2.1 Simulation framework

#### 2.1.1 Local adaptation in a continuous landscape

We simulated local adaptation in a continuous landscape using genetically explicit, forward-in-time simulations implemented in Nemo (v2.4a) (Guillaume and Rougemont, 2006) in a Write-Fisher frame-work with migration, selection and promiscuous random mating. The landscape consisted of a 48 × 48 grid of demes, each with carrying capacity K_d_ = 5 individuals (K_tot_ = 11,520). Spatially autocorrelated environmental optima were generated in R using the gstat package (Pebesma, 2004), assuming an exponential variogram (sill = 10, range = 50, nugget = 0.01). For a given landscape map (Fig. S1), a random field was generated and rescaled to [0, 1].

Dispersal followed an isotropic Gaussian kernel (Nathan et al., 2008),

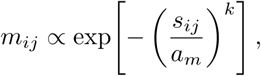

where where *s_ij_* denotes the Euclidean spatial distance between demes *i* and *j*, *a_m_* = 2 × 10^−5^ controls mean dispersal distance and *k* ∈ {1, 1.5, 2} defines kernel shape, corresponding to high, medium, and low gene flow. Migration probabilities were normalized to sum to one, and absorbing boundaries were assumed at grid edges.

#### 2.1.2 Recombination map construction

Recombination maps were derived from empirical nine-spined stickleback Marey maps for chromosomes 1-10 (Kivikoski et al., 2021). Local recombination rates were calculated as the mean of male- and female-specific estimates, yielding chromosome lengths of 73–120 cM across 15–33 Mbp. To increase the probability that neutral SNPs fall into measurable LD with causal loci, each map was scaled to one tenth of its empirical length.

For each replicate, two chromosome types were simulated: a *QTN* chromosome containing both quantitative trait nucleotides (QTNs) and neutral loci, and a *ntrl* chromosome containing only neutral loci. On *QTN* chromosomes, 100 biallelic QTNs were placed at random map positions. Allelic effects *β* were drawn from an exponential distribution (mean ≈ 0.05), with opposing effects (±*β*) at each locus. Individual genotypic values were calculated as

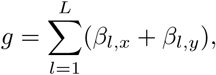

where *β_l,x_* and *β_l,y_* denote maternally and paternally inherited effects at locus *l*.

To increase resolution around QTNs, 10^5^ neutral SNPs were sampled from the recombination map with elevated density near QTN positions; neutral SNP locations were generated by inverse transform sampling from a kernel density estimate of QTN positions, ensuring strong spatial correspondence between neutral and causal loci (*r*^2^ *>* 0.6). Neutral chromosomes used identical SNP positions, but all QTN effect sizes were set to zero.

#### 2.1.3 Trait architecture and fitness landscape

A single quantitative trait was simulated under Gaussian stabilizing selection around the local environmental optimum. Individual fitness was defined as

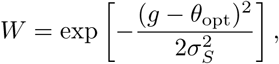

where *θ*_opt_ is the local optimum and *σ*^2^ ∈ {0.5, 1, 2} controls selection strength. Fitness directly deter-mined offspring survival, with exactly K_d_ surviving offspring per deme each generation.

Simulations included a burn-in of 10^4^ non-overlapping generations with a uniform optimum (*θ*_opt_ = 1), followed by 10^3^ generations local adaption to the selection landscapes. Loci started out alternatively homozygote for −*β* and +*β* to ensure *g* ≈ 0 at start-up. During burn-in, genetic variation was gener-ated by mutation at both neutral sites and QTNs as the population adapted to a mean *g* ≈ 1 resulting in realistic linkage disequilibrium relationships between QTNs and neutral SNPs.

Neutral loci mutated at rate *µ* = 5 × 10^−7^, yielding ∼ 1500 SNPs per chromosome with minor allele frequency maf *>* 0.05 at the end of simulations. QTNs mutated at *µ* = 5 × 10^−8^ to maintain identity by descent. All combinations of dispersal regime and selection strength were simulated across ten replicate landscape maps (Fig. S1) and ten recombination maps, yielding 900 simulations. We uniformly sam-pled 125 spatial coordinates from the 48 × 48 deme grid and sampled one individual per location for all downstream analyses. These coordinates were fixed and reused across all landscape replicates to ensure that differences among replicates reflect variation in selection and demography rather than stochastic differences in spatial sampling.

The ratio *Q_ST_ /F_ST_* was used to quantify the strength of spatially divergent selection, providing a stan-dardized measure of selection efficacy across simulation scenarios (Leinonen et al., 2013) and the level of local adaptation was estimated by the correlation between individual phenotypic values (*g*) and their local optima (*θ*_opt_).

### 2.2 Estimation of LD decay

All pairwise LD values were estimated as *r*^2^ using SNPRelate::snpgdsLDMat (Zheng et al., 2012). LD decay was quantified by fitting

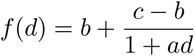

to the 95th percentile of *r*^2^ values across non-overlapping 5 kb windows (Vos et al., 2017) using non-linear least square regression (R-function nls; algorithm=“port”; Bates and Watts, 1988). Here, *b* denotes background LD, *a* the decay rate, and *c* the intercept at *d* = 1, where *d* denotes physical distance in bps between loci along the chromosome. For empirical data, decay parameters were estimated in sliding 1 Mbp windows and summarized by their median. LD-based thresholds were expressed relative to the empirical decay curve using the quantile parameter *ρ*, where

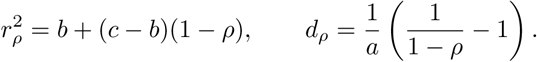

This normalization allows direct comparison across datasets with different LD landscapes. On this rel-ative scale, *ρ* = 1 corresponds to the point where LD has decayed to the background level, and *ρ <* 1 represents increasing *r*^2^’s and decreasing *d*’s (between pairs of loci). Parameters without *ρ* subscript, refer to the same quantity on the absolute scale or to the parameter in conceptual terms.

### 2.3 Genotype by environment association analyses

We tested genotype by environment associations using latent factor mixed models (LFMM) using R-package LEA (Frichot and Fraņcois, 2015) and linear mixed models using EMMAX (Kang et al., 2010). For LFMM, the number of latent factors *K* was inferred by clustering the first ten principal compo-nents of the genotype matrix using factoextra::eclust (Kassambara and Mundt, 2020). LFMM models were fitted using lfmm2 with genomic control enabled.

EMMAX analyses were performed using published R implementations (Li et al., 2018; Fang et al., 2021). Genetic relationship matrices (SNPRelate::GRM) were estimated from LD-pruned SNP sets as recommended by Lotterhos (2019) . LD-pruning was achieved by single linkage clustering for pairwise LD values estimated in 1000 bps windows using R-package igraph (Cśardi and Nepusz, 2006; Antonov et al., 2023). An edge list only containing rows with *r*^2^ *> r*^2^ (the half way decay distance) was converted into a *graph object* (graph from edgelist; directed=FALSE) and decomposed into sub-graphs (decompose) retaining one randomly selected SNP from each. Genomic control (GC; (Devlin and Roeder, 2004; Price et al., 2010)) was used to correct residual p-value inflation by normalizing test statistics to the expected null median.

Outlier analyses were performed separately for each simulated data set but chromosomes simulated under different recombination maps but with identical selection landscapes were concatenated into a single artificial genome of 20 chromosomes (10 *QTN*, 10 *ntrl* ), prior to FDR corrections (p.adjust, method = “fdr” in R) to better reflect genome-scale multiple testing. Because the sampling coordinates were fixed for all simulations (see above), the aligned genotype–environment relationships were preserved mimicking independently segregating chromosomes experiencing identical selection surfaces. This substantially increased the number of segregating causal variants (true positives) available for evaluating outlier detection performance. Chromosomes were labeled 1–20, with odd-numbered chro-mosomes containing QTNs.

### 2.4 Combining association statistics with local linkage disequilibrium

To incorporate local LD information, we combined association test statistics *F* from EMMAX and LFMM with local LD summary statistics *ld_w_*, defined as the median *r*^2^ within a window of size *w*. Let *F*_org_ denote the original association statistic, and define the observed normalized LD-weighted statistic as

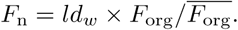

An empirical null distribution, *F*_p_, was obtained by circularly permuting (Cabrera et al., 2012) the *ld_w_* vector prior to multiplication with *F*_org_*/F*_org_. This procedure preserves the spatial autocorrelation structure of LD while randomizing its association with the *F* test statistic.

In a quantile–quantile plot using the theoretical *F* distribution (*F*_0_) as the baseline, deviations of the permuted statistic *Q*(*F*_p_) from *Q*(*F*_0_) (grey shaded area in Fig. 1) quantify the systematic distortion that arises when association statistics are multiplied by local LD estimates. This distortion is expected under the null hypothesis (*H*_0_) that *ld_w_* is unrelated to selection, because the product of two right-skewed variables produces heavier-tailed distributions even in the absence of true signal. In contrast, when local LD is elevated due to physical proximity to QTNs, then the quantiles of the observed LD-weighted statistic *Q*(*F*_n_) are expected to exceed *Q*(*F*_p_) (orange shaded area in Fig. 1) representing a true signal of selection.

**Figure 1:**
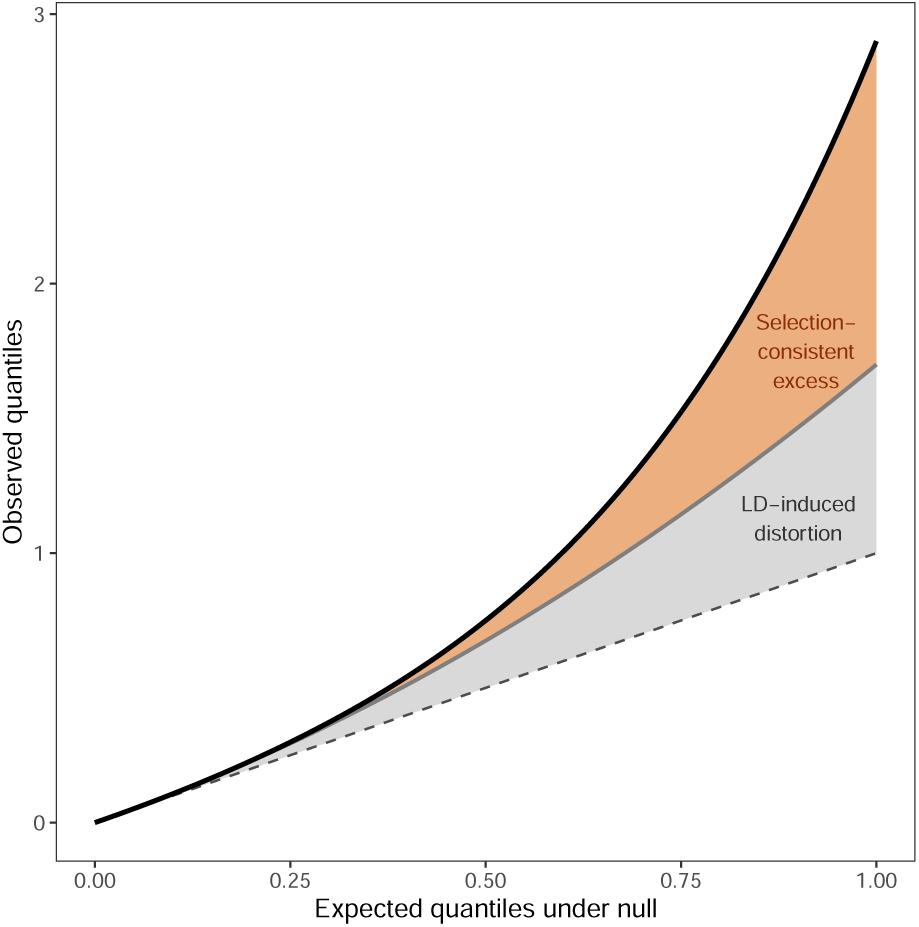
Conceptual illustration of LD-scaling and quantile transformation of association statistics. The dashed line represents the theoretical null expectation (*F*_0_). Weighting association statistics by local linkage disequilibrium (LD) introduces systematic deviations from the null (grey curve and shaded area, *F_p_*), even under neutrality. The observed LD-weighted statistic (*F_n_*, black curve) combines this LD-induced distortion with excess signal consistent with selection (orange shaded area). The quantile transformation corrects for the LD-induced component while preserving relative ranking in the upper tail, enabling calibrated inference under complex LD structure. An example from empirical data is shown in Figure. S2.

To correct for LD-induced distortion, we adjusted *Q*(*F*_n_) by the difference between *Q*(*F*_p_) and *Q*(*F*_0_), yielding the quantile-transformed statistic *F*_q_ (Fig. S2). Under the null hypothesis *H*_0_ that local LD is unrelated to the probability that a SNP is associated with selection, this transformation ensures *Q*(*F*_q_) ≈ *Q*(*F*_0_), allowing p-values to be derived directly from the theoretical *F* distribution with appropriate degrees of freedom.

In practice, when *Q*(*F*_p_) increases more rapidly than *Q*(*F*_n_), the *F*_q_ transformation may locally distort SNP ranking (Fig. S2). To preserve correct ordering in the upper tail, we therefore define the final test statistic as

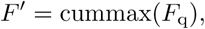

which effectively truncates the distribution in regions where *Q*(*F*_p_) overtakes *Q*(*F*_n_).

Because linear interpolation is used (approx function in R), the resulting *F* ^′^ statistics were nearly identical across independent permutations of the *F*_p_ vectors (*r*^2^ *>* 0.99 ). We nevertheless used ten replicate permutations throughout the study. No genomic control was required for *F* ^′^, as the inflation factor was consistently ≪ 1.

### 2.5 Using outlier regions (ORs) as independent test units

Traditionally the performance of outlier detection methods has been evaluated by treating individual SNPs as independent test units which greatly deflates recall when multiple linked SNPs are in LD with the same causal variant. We thus evaluated our method’s performance at the level of *outlier regions* (ORs) representing clusters of physically linked and correlated outlier SNPs putatively associated with distinct genomic signals of selection.

For each simulated dataset, a *graph object* was constructed from an edge list of all outlier loci (for a given significance threshold *α*), containing only within chromosome SNP pairs with *r*^2^ *> τ*_LD*,ρ*_ and physical distance *d < τ*_d*,ρ*_, where *ρ* denotes the quantile position along the empirical LD-decay curve. The resulting graph was then decomposed into subgraphs, each representing an OR (Fig. 2). The parameters *τ*_LD*,ρ*_ and *τ*_d*,ρ*_ thus define how strongly associated and how physically close two SNPs must be to be assigned to the same OR.

**Figure 2:**
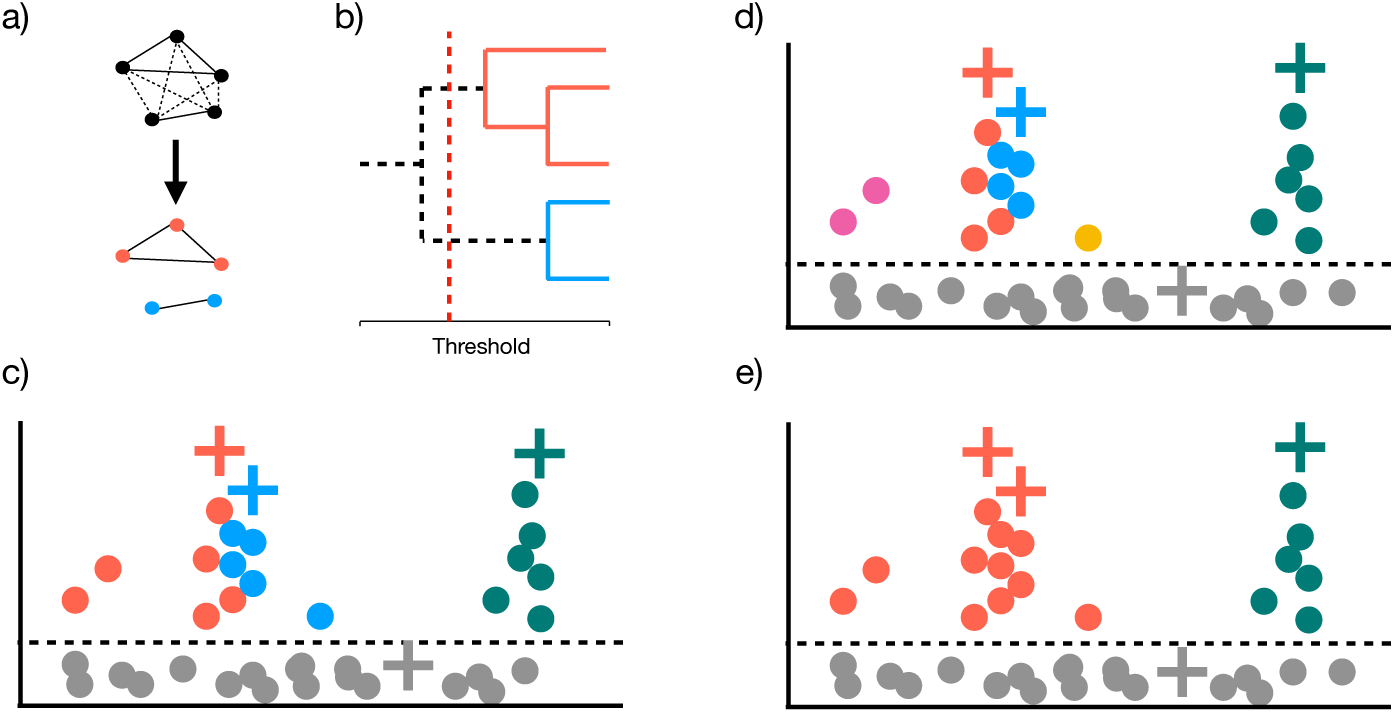
Single-linkage clustering of outlier loci. (a–b) Schematic overview of the single-linkage clustering procedure. Edges connecting outlier SNPs with *r*^2^ *< τ*_LD_ or *d > τ*_d_ (dashed lines) are removed, and the remaining connected components define outlier regions (ORs), shown in distinct colors. Panel (a) shows the network representation, and (b) the corresponding single-linkage clustering tree with the applied threshold indicated by a red vertical line. (c) Ideally, each OR corresponds one-to-one with a quantitative trait nucleotide (QTN, indicated by “+”), but inappropriate threshold choices can distort this relationship (d–e). (d) High *τ*_LD_ and low *τ*_d_ values lead to *over-clustering*, where multiple ORs correspond to the same QTN (purple and yellow represent ”satellite” ORs), whereas in (e) low *τ*_LD_ and high *τ*_d_ values cause *over-merging*, where several QTNs fall within a single OR. In (c–e), true positives (TP) are distinct QTNs captured by separate ORs; false positives (FP) are the total number of ORs minus TP; and false negatives (FN) are QTNs that (i) do not exceed the significance threshold and (ii) are not associated with any outlier locus by *r*^2^ *> τ*_LD*,ρ*=0.75_ or *d > τ*_d*,ρ*=0.95_ (grey “+” symbols). The corresponding precision–recall (PR) scores for panels (c–e) are 0.75 and 0.375 for both (d) and (e), since in (d) a single QTN_f_ per OR is considered. Note that ORs can also be defined jointly by considering outliers detected by any of a given set of methods.

True positive QTNs were defined as causal SNPs that contributed at least 5% of the total additive genetic variance among QTNs with maf *>* 0.05 on the same *QTN* chromosome. The proportional contribution of each QTN was calculated as

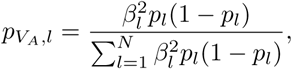

where *p_l_* is the frequency of the reference allele and *N* is the number of segregating QTNs per chromosome (maf*>*0.05) (Lotterhos, 2019).

For OR, we assigned a single focal QTN (QTN_f_) by identifying the causal variant in highest LD with any SNP in the region, provided the association exceeded *r*^2^ *> r*^2^ and the physical distance was closer than *d < d_ρ_*_=0.95_. When multiple candidate QTNs met these criteria, we selected the one with the highest LD with any of the neutral SNPs in the OR, and duplicate assignments were removed so that each true QTN could be counted at most once.

An OR was classified as a true positive (TP) if its assigned QTN_f_ was among the true positive QTNs. ORs that did not map to any such QTN_f_ were classified as false positives (FP), and false negatives (FN) were defined as true positive QTNs not captured by any OR. Precision, recall, and their product (PR) were then computed as:

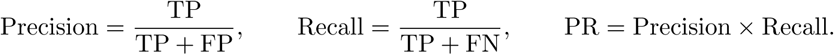

Ideally, each QTN_f_ should be captured by exactly one OR, and no OR should contain multiple QTN_f_’s (Fig. 2b). Both poor performance and suboptimal clustering result in deviations from this one-to-one correspondence. Too high *τ*_LD_ or too low *τ_d_* results in *over-clustering*, generating multiple small ORs (“satellite clusters”) corresponding to the same QTN_f_ and are thus by definition false positives (Fig. 2c).

In contrast, too low *τ*_LD_ or too high *τ_d_*results in *over-merging* where multiple distinct QTNs belong to a single OR (Fig. 2d) thus underestimating the number of true positives. We also introduced an alternative and complementary way to reduce satellite clusters and false positive ORs caused by genetic structuring by filtering out OR with fewer than *l_min_* loci (Li et al., 2018; Fang et al., 2021).

### 2.6 Accounting for uncertainty in parameter choice using AUC-PR^∗^

Previously, the performance of outlier methods have mainly been evalutaed using precision–recall (PR) metrics, providing an overall measure of method performance that is independent of any single significance threshold (*α*) (e.g., Lotterhos et al., 2022). To account for uncertainty in the additional parameter choices introduced here, we extended the standard AUC-PR into a multidimensional framework, denoted AUC-PR^∗^ that summarizes cumulative PR performance across random draws of key parameters (*w*, *α*, *τ*_LD_, *τ*_d_, and *l*_min_). Based on pilot runs exploring the parameter space, each random draw represents a combination of parameter settings sampled uniformly from the following ranges:

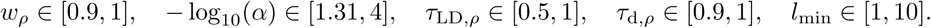

Because *ld_w_*-values for *F* ^′^-estimation were the most computationally expensive to calculate, we first generated *ld_w_*-vectors for 200 random draws of *w_ρ_*, and for each 25 additional draws were generated for the remaining threshold parameters, resulting in 5,000 parameter combinations in total per method and simulation scenario. Only *w* is intrinsic to the LD-weighted statistic *F* ^′^, while the remaining parameters define significance and clustering thresholds to distinguish true and false ORs. For each method, cumulative PR curves were constructed by taking the running maximum of PR values across parameter draws:

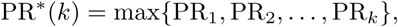

where *k* indexes the number of random draws. Methods for which PR^∗^(*k*) rises steeply and reaches a high asymptote achieve larger AUC-PR^∗^ values, reflecting greater power and robustness. The AUC was calculated by integrating the mean PR^∗^ curves across 500 bootstrap replicates of 500 random parameter subsets (using the trapezoidal rule in R).

To facilitate comparison among methods, we normalized the area under the precision–recall curve (AUC-PR), irrespective of how it was constructed, by its empirical maximum max(PR). This yields the ratio

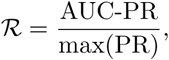

which provides a standardized, unitless measure of robustness that is directly comparable across methods and performance metrics.

### 2.7 Consistency scoring and dimensionality reduction

Because true outlier loci tend to cluster near their QTN_f_’s, SNPs genuinely associated with causal variants are expected to be detected as outliers more consistently across parameter settings than false positives. To leverage this we integrated over parameter uncertainty by calculating a consistency score (*C*) for each SNP, defined as the proportion of random parameter combinations (draws) in which that SNP was included in an OR (Fang et al., 2021).

To evaluate the performance of the *C*-score in differentiating true from false positive ORs, we adapted the AUC-PR^∗^ framework by replacing the three parameters most sensitive to uncertainty–the LD window size (*w*), the outlier significance threshold (*α*), and the minimum OR size (*l*_min_)–with a single threshold on the consistency score, *τ*_C_ ∈ [0, 0.5]. The remaining clustering parameters (*τ*_LD_*, τ*_d_), which determine how outlier SNPs are grouped into ORs, were retained. We then performed 5,000 random draws across the full parameter set (*τ*_C_*, τ*_LD_*, τ*_d_), computing precision and recall at each draw as in the standard AUC-PR^∗^ procedure, denoted AUC-PR_C_.

In addition to considering each of the four ”base” methods–EMMAX (EMX), LD-scaled EMMAX (EMX^′^), LFMM (LFMM), and LD-scaled LFMM (LFMM^′^)–independently, we also evaluated all the methods jointly (Joint) by combining SNPs identified as outliers by any of the four methods before defining ORs.

Influence of simulation parameters and method choice on simulation outputs and performance metrics were estimated using partial *η*^2^ from Type-III ANOVA models (R package effectsize). To identify parameter combinations associated with optimal performance, for each simulation replicate × method × scenario, we ranked parameter draws by PR and retained the top 10% of the draws with ties broken at random. This provides a direct summary of the regions in parameter space that yield the best trade-off between precision and recall, independent of the cumulative performance metric AUC-PR^∗^. Co-dependencies among threshold parameters were assessed using logistic regression, classifying parameter draws in the top 10% of PR values and testing pairwise interactions via likelihood-ratio tests. The strength of co-dependency was quantified by the improvement in model fit when including the interaction, using a likelihood ratio test (*χ*^2^) and the corresponding change in Akaike’s Information Criterion (Δ*AIC* = *AIC*_reduced_ − *AIC*_full_).

### 2.8 Re-analysis of empirical stickleback data

A common strategy for separating genome-wide signals driven by drift and demographic history from locus-specific signals of selection is to analyse systems in which similar adaptations have evolved repeatedly in multiple populations derived from a shared ancestral background (e.g. Johannesson, 2001; Schluter et al., 2004). The three- and nine-spined sticklebacks (*Gasterosteus aculeatus* and *Pungitius pungitius*) provide a well-established comparative framework for this purpose because both repeatedly form marine and freshwater ecotypes across the northern hemisphere, yet differ in population connectivity and background genetic differentiation (McKinnon and Rundle, 2002; Foster and Baker, 2004; DeFaveri et al., 2012; Guo et al., 2013; Fang et al., 2021; Kemppainen et al., 2021). We therefore re-analysed empirical data set from North-Eastern Atlantic populations Fang et al. (2021), for which LDna results are available, to benchmark the LD-scaled statistic (*F* ^′^) and the outlier-region (OR) framework under realistic LD structure.

We used the phenotype data and pre-filtered genotype-likelihood data released by Fang et al. (2021) (three-spined: 881,786 SNPs from 117 individuals; nine-spined: 1,344,316 SNPs from 145 individuals; Zenodo accession: 4722879). Expected genotypes were used for association testing with EMMAX and LFMM, whereas called genotypes were used for LD-based analyses implemented in SNPRelate.

Empirical analyses followed the same workflow as for the simulated data: (i) estimation of chromosome-specific LD-decay curves; (ii) LFMM and EMMAX association testing (iii) generation of LD-window scaling vectors *ld_w,ρ_* by drawing *ρ* ∼ Unif(0.75, 1) (200 draws); and (iv) for each *ld_w,ρ_* draw, sampling 25 parameter combinations for *α*, *τ*_LD_, and *τ*_d_ from the same uniform ranges used in the simulations. However, the upper bound of *l*_min_ was adjusted to *l*_min_ = 30 and *l*_min_ = 45 for three- and nine-spined sticklebacks, respectively, to reflect the higher SNP densities per cM. In addition, for nine-spined sticklebacks, lineage (Eastern, Western, admixed) was additionally included as a fixed-effect covariate in EMMAX analyses, following Fang et al. (2021). The resulting parameter draws across all methods were used to compute SNP-level *C*-scores for Joint_C_ and compared to the FDR-corrected p-values from LFMM, enabling direct comparison of inference sensitivity to local LD structure in the empirical data. All analyses were implemented in R using custom code (see Data and Code Availability).

## 3 Results

### 3.1 Drivers of local adaptation signals and implications for detectability

Across simulation scenarios, variation in genomic and phenotypic structure was dominated by gene flow. Both genetic differentiation (*F_ST_* ; 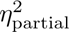= 1.00) and phenotypic differentiation (*Q_ST_* ; 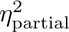= 0.801) decreased with increasing gene flow, whereas the ratio *Q_ST_ /F_ST_* increased (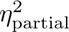 = 0.916), reflecting the need for stronger or more heterogeneous selection to maintain phenotypic divergence under high migration (Fig. 3; Fig. S3). Background LD (*b*) also increased with gene flow (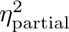 = 0.873), whereas selection intensity had comparatively weak effects on these global summaries (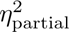 *<* 0.2). Other simulation outputs showed limited sensitivity to demographic parameters (Fig. S3). LD-decay rates (*a*), the distribution of allelic effect sizes among segregating loci, and the proportion of additive genetic variance explained by individual QTNs were largely invariant across scenarios (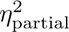 *<* 0.1). Nevertheless, LD decayed more slowly on QTN-bearing chromosomes than on neutral chromosomes (mean *a*_QTN_*/a*_ntrl_ = 0.84; Fig. 3), consistent with local inflation of LD around selected loci. Together, these results confirm that the simulations generated biologically coherent gradients of adaptive divergence and LD structure, capturing key evolutionary regimes relevant to genome scans in natural popu-lations.

**Figure 3:**
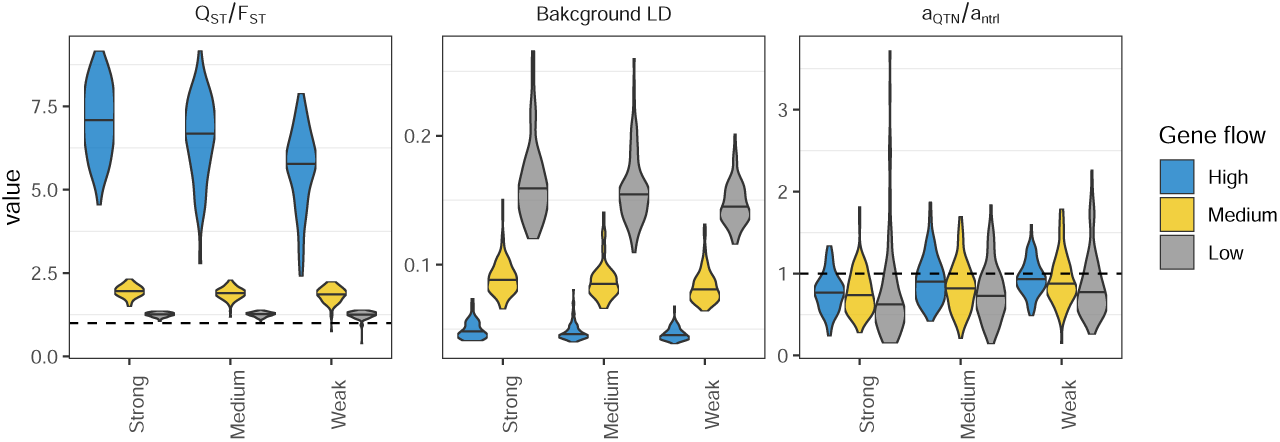
Key simulation summary statistics. Shows three summary statistics; *Q_ST_ /F_ST_* , background LD (*b*) and the ratio between LD-decay rate (*a*) between chromosomes with QTN and neutral chromo-somes. Horizontal dashed lines indicate where *Q*_ST_ = *F*_ST_ and *a*_QTN_ = *a*_ntrl_. contexts.

### 3.2 Robustness of parameter choice

Across the top 10% of parameter draws ranked by precision–recall, summed partial *η*^2^ values were uniformly small (*η*^2^ *<* 0.1), indicating that parameter regions yielding high performance were largely invariant across demographic scenarios and inference methods (Fig. 4a). In particular, parameters expressed relative to LD decay (*w_ρ_*, *τ*_LD*,ρ*_, *τ*_d*,ρ*_) showed minimal sensitivity to gene flow or selection intensity (*η*^2^ *<* 0.02). This stability supports the use of LD-scaled thresholds as a means of reducing dependence on dataset-specific demographic properties, thereby improving generality across evolutionary

**Figure 4:**
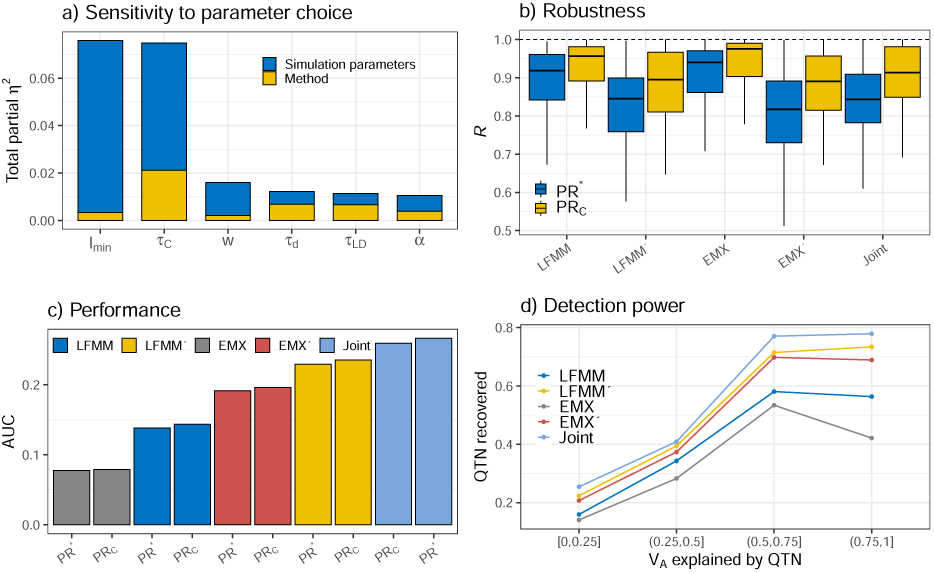
Summary of parameter sensitivity and method performance. (a) Partitioning of effect sizes (*η*^2^) among the 10% highest-performing parameter draws (out of 5,000) shows that statistical power (PR) is not sensitive to parameter choice, each parameter explaining *<* 10% of the variation in PR. (b) The ratio Summary of performance metrics from the top R = AUC-PR^∗^*/* max(PR) quantifies how close each method’s average performance across draws is to the maximum among all draws shown for EM-MAX/LFMM analyses before and after (indicated by the ”′”) scaling the *F* -values with LD. Replacing three of the parameters (*α*, *l*_min_ and *w*) with the composite consistency score (*C*) increases robustness, as indicated by higher R values (data pooled across all simulations). (c) Overall performance (AUC ) of outlier detection methods using the full set of parameters (PR^∗^) or the reduced set of parameters (PR_C_), pooled across all simulations (see Fig. 6 for details). (d) Statistical power as a function of the proportion of *V_A_*explained by a QTN. In Joint analyses, outliers from all methods (EMMAX/LFMM with scaled and non-scaled *F* -values) were considered when defining outlier regions. expectations and demonstrate that LD-scaled, consistency-based integration yields fewer but more co-herent and biologically interpretable candidate regions.

### 3.3 Method performance

Across all scenarios and methods, robustness (measured as R = AUC-PR*/* max(PR)) increased when using the *C*-score (Fig. 4b) but results based on AUC-PR^∗^ and AUC-PR_C_ were almost identical (*η*^2^ *<* 0.001; Fig. 4b; Fig. S4). This indicates that while a higher maximum performance can be potentially achieved by using the full set of parameters, the number of parameter combinations that yields close to maximum performance are expected to be less common among draws than in consistency-based analyses.

An illustrative example of how consistency-based inference improves localization of causal variants is shown in Fig. 5a–d. Compared to baseline LFMM results (Fig. 5a), Joint_C_ more clearly differentiates genomic regions associated with QTNs, reflecting its close association with locally elevated LD between SNPs and their focal causal variants (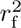). Consistent with this interpretation, Joint_C_ values show a stronger correlation with 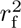 than FDR-corrected LFMM *q*-values (Fig. 5c), explaining approximately twice as much variation (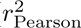 = 0.44 versus 0.23). This amplification of signal arises because Joint_C_ integrates LD information across parameter draws, thereby prioritizing SNPs embedded within distinct LD signatures around QTNs. Importantly, local LD itself is non-randomly associated with selection in this example, as indicated by a positive correlation between *ld_w_* and 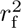 (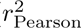 = 0.14; Fig. 5d). These patterns demonstrate that leveraging LD structure through consistency-based aggregation sub-stantially improves the contrast between true selection signals and background variation relative to SNP-level inference alone.

**Figure 5:**
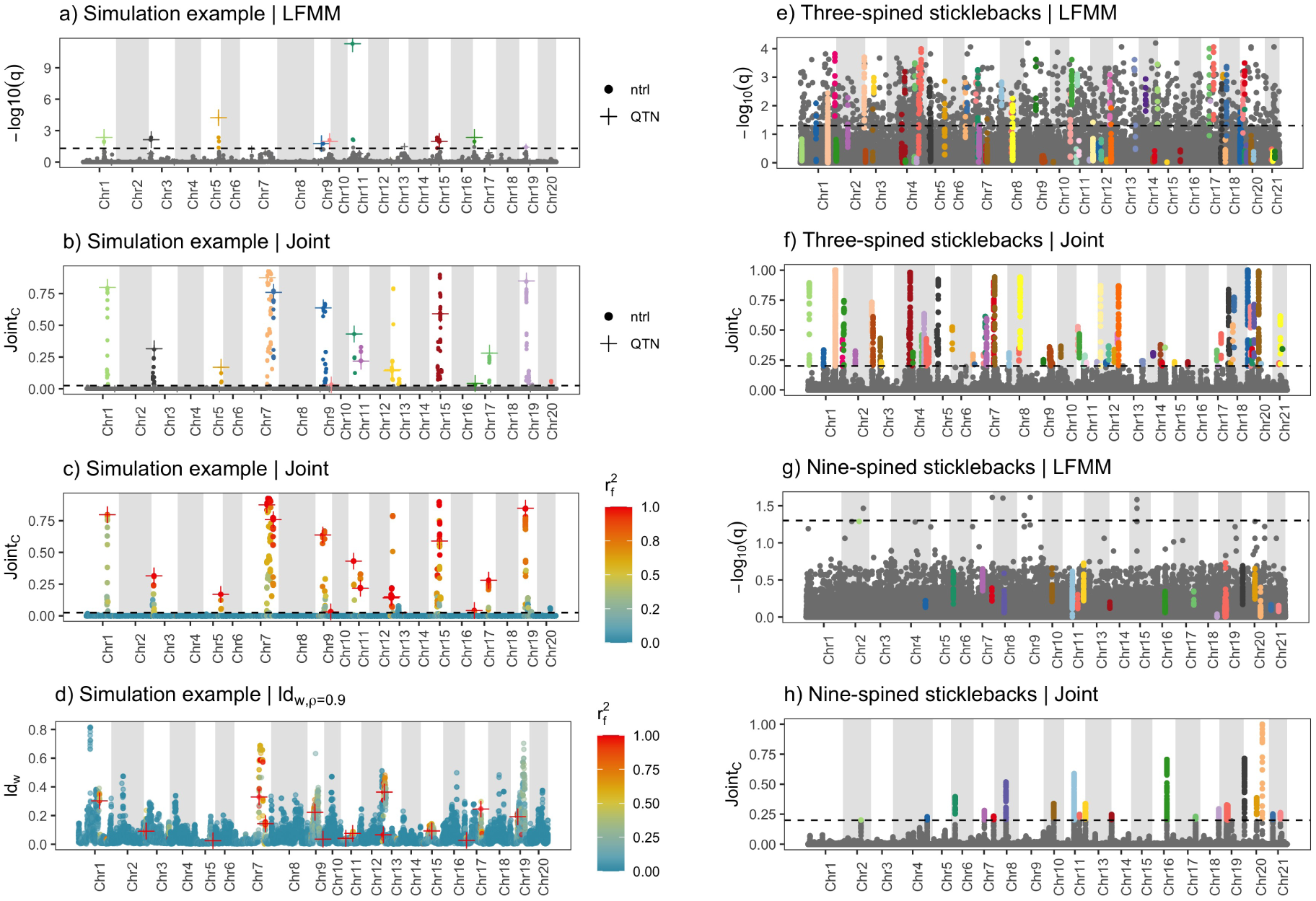
Comparison of LFMM and joint outlier analyses in simulated and empirical data. Panels (a–d) show example outlier results from the highest-performing random draw in a simulated genome under high *gene flow* and strong *selection*. Results are shown for LFMM (a; TP = 8, FP = 0, FN = 17, PR = 0.32) and for joint analyses aggregating evidence across all base methods (Joint_C_; b–d; TP = 14, FP = 1, FN = 11, PR = 0.52). Colors denote matching outlier regions (ORs), de-fined using *τ*_LD*,ρ*_ = 0.75 and *τ*_d*,ρ*_ = 0.99 for LFMM, and *τ*_LD*,ρ*_ = 0.99 and *τ*_d*,ρ*_ = 0.99 for Joint_C_. ORs sharing the same color across panels correspond to the same genomic region; causal loci are indi-cated by “+” and neutral loci by “·”. Panel (c) is identical to (b) but colored by the strength of asso-ciation between SNPs and their focal causal locus (r^2^). Panel (d) shows the same SNPs as in (c), but with *ld_w_*_=0.9_ on the *y*-axis instead of Joint_C_. Panels (e–h) show re-analyses of empirical three-spined (e,f) and nine-spined (g,h) stickleback data using LFMM (e,g) and joint analyses (f,h). Colors repre-sent distinct ORs identified by Joint_C_ using *τ*_LD*,ρ*_ = *τ*_d*,ρ*_ = 0.999, yielding 49 and 20 ORs in three- and nine-spined sticklebacks, respectively. The dashed horizontal line in joint analyses denotes the Joint_C_ threshold *τ*_C_ = 0.2. For LFMM analyses (a,e,g), dashed horizontal lines indicate the significance thresh-old for FDR-corrected *p*-values (− log_10_ *q*) at *α* = 0.05.

Because the consistency score *C* integrates over uncertainty in three of the five tuning parameters (*w*, *α*, and *l*_min_), defining outliers are reduced to the choice of a single threshold, *τ*_C_, effectively replacing the nominal significance threshold *α*. *C*-based analyses are therefore particularly relevant for empirical data where the lack of ground truth prevents systematic optimization of the full parameter set. Subsequent comparisons are therefore based on AUC-PR_C_, using LFMM_C_ as a baseline. Across all simulation scenarios, incorporating LD information improved performance for both association methods. Baseline EMMAX had the lowest mean performance (AUC-PR_C_ = 0.079), but LD-scaling increased performance by 150% (AUC-PR_C_ = 0.196), exceeding baseline LFMM by 37%. LD-scaling also substantially improved LFMM performance (AUC-PR_C_ = 0.235, +65%). Consequently, differences between EMMAX and LFMM narrowed once local LD structure was explicitly leveraged (Fig. 4c).

Aggregating evidence across methods before defining outlier regions (Joint) yielded the highest overall performance (AUC-PR_C_ = 0.260), representing an 81.2% improvement over baseline LFMM. Detection power increased with the per-locus proportion of additive genetic variance explained, *p_VA_,i* (Fig. 4d), and the performance differences among methods were consistent with broadly increased power across the full range of *p_VA_,l* rather than being confined to a specific range of values.

Across simulation scenarios and replicates (Fig. 6a), Joint_C_ was the best-performing approach (or within 5% of the best) in 66% of analyses, followed by LFMM^′^ (48%) and EMX^′^ (22%). Importantly, LD-scaled and consistency-based methods were substantially less likely to fail completely under low gene flow (6.7% of the replicates), where baseline LFMM failed to detect any true positives in ∼2/3 of the cases.

**Figure 6:**
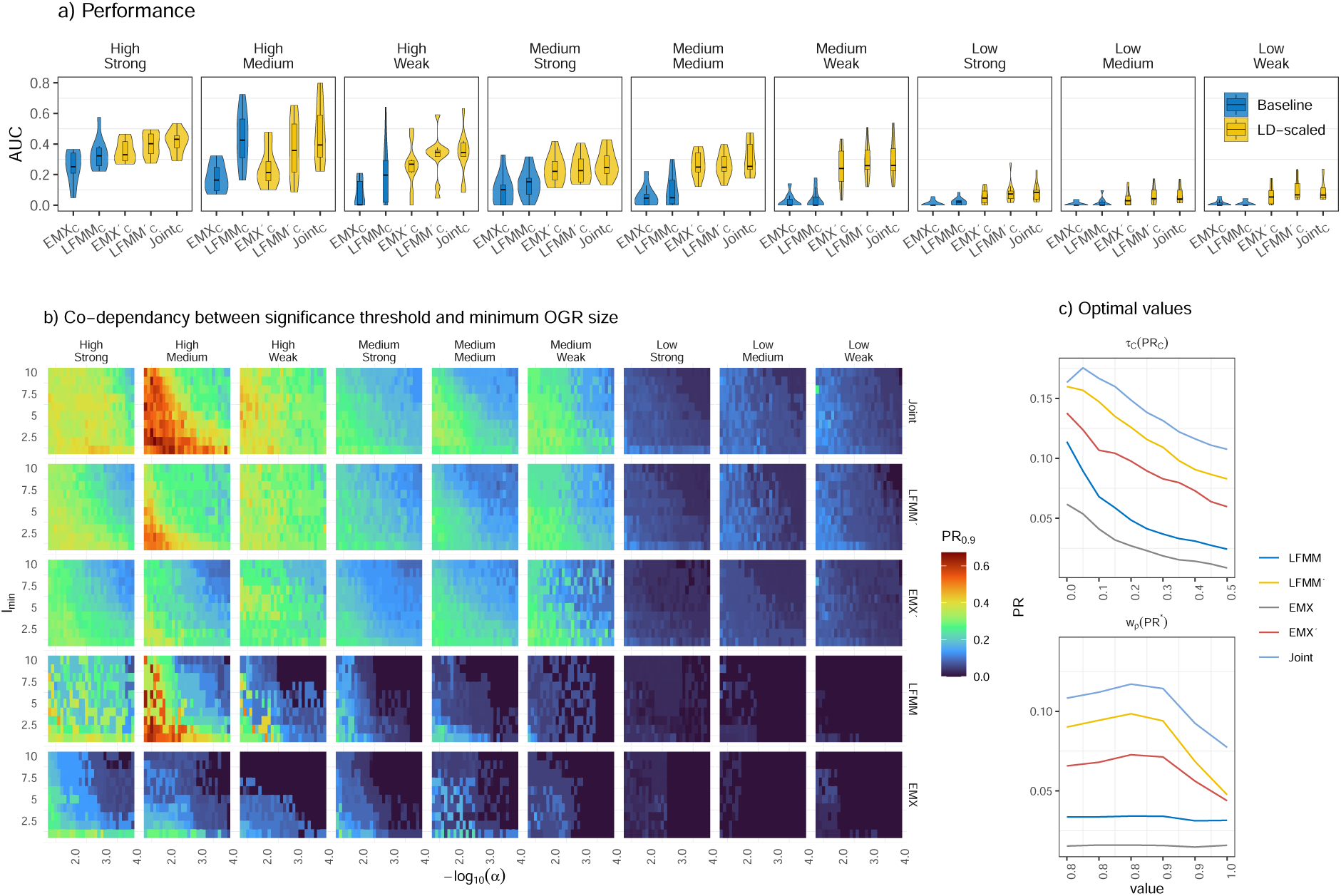
Performance of outlier detection methods across simulation scenarios. (a) Shows method performance based on AUC-PR_C_ calculated from 5,000 random parameter draws of *τ*_C_, *τ*_LD_ and *τ*_d_. Joint indicates that outliers across all methods were considered when partitioning them into ORs. Panels right to left represent simulation scenarios in order of decreasing *gene flow* (high, medium, low) and *selection intensity* (strong, medium, weak). Methods are in order of increasing mean performance across all simulation scenarios (left to right). Color distinguishes methods based on original *F* -values (blue) and those that rely on LD-scaled values (yellow). (b) Heat maps represents binned 90% quantile of PR for different parameter combinations of significance threshold (*α*) and minimum OR size (*l*_min_) illustrating their co-dependency. Co-dependencies for other parameter combinations are shown in Supplementary Fig. S5b-e. (c) Shows moving average of PR across the 5,000 parameter as a function of *τ*_C_ (used for AUC-PR_C_) and *w_ρ_* used to calculate *ld_w_* (used for AUC-PR^∗^). Base method is indicated by color as shown in legend.

Despite the same recombination landscape in *QTN* and *ntrl* chromosomes within a simulation replicate, false positive rates were consistently higher on *QTN* chromosomes than on *ntrl* chromosomes, except under low gene flow, where the mean number of false positive ORs increased sharply on *ntrl* chromosomes approaching those observed on *QTN* chromosomes (Fig. S4).

### 3.4 Co-dependencies among parameters

Although increasing threshold parameters (− log_10_ *α*, *τ*_C_, *l*_min_, *τ*_LD_, *τ*_d_) generally reduced false positives (Fig. S6), heatmaps of the upper tail of PR revealed strong co-dependencies among thresholds (Fig. 6b; Fig. S5). The most consistent interaction occurred between the significance threshold *α* and the minimum OR size *l*_min_, where increased stringency in one parameter could be compensated by relaxation of the other. This demonstrates that OR size can be as reliable at predicting false positives as the effect size of the association itself (proportional to test significance). In contrast, co-dependencies involving *τ*_C_ were weaker, reflecting its role as an integrative summary of multiple sources of uncertainty.

Despite these interactions, the shape of high-performance regions in parameter space was highly concordant across methods and simulation scenarios (Fig. 6b; Fig. S5). For example, optimal performance consistently occurred at *w_ρ_* ≈ 0.8–0.9 across all methods (Fig. 6c).

However, among the top 10% parameter draws *τ*_C_, *α* and *l*_min_ exhibited substantial variability, with wide interquartile ranges (*Q*_25_–*Q*_75_: 0.049-0.238, 0.0028-0.035 and 1–5 for *τ*_C_, *α* and *l*_min_, respectively). Thus, while less stringent values often produced the highest performances, a broad range of higher thresholds also yielded near-optimal trade-offs between true and false positive rates across these parameters. Together, these patterns demonstrate that no single threshold value is universally optimal, motivating the use of conservative parameter choices within empirically supported ranges for analyses of empirical data.

### 3.5 Re-analyses of empirical data

Background LD levels in three-spined sticklebacks (*b* = 0.050) closely matched those observed in simulations with high gene flow, whereas background LD in nine-spined sticklebacks (*b* = 0.20) exceeded all simulated scenarios, consistent with stronger genome-wide differentiation. LD-decay rates differed more between the two species (1.88×) than between low and high gene flow scenarios in the simulations (1.2×), emphasizing the importance of relative LD scaling.

Baseline LFMM analyses showed limited utility in both species: extensive background signal in three-spined sticklebacks and low statistical power in nine-spined sticklebacks (Fig. 4e,g). In contrast, the joint consistency-based approach (Joint_C_) identified 49 and 20 outlier regions in three- and nine-spined sticklebacks, respectively, using *τ*_C_ = 0.2 and stringent clustering thresholds (*τ_LD,ρ_* = *τ_d,ρ_* = 0.999). This *τ*_C_ value lies near the upper quartile of values observed among top-performing simulation draws and was chosen to prioritize robustness over maximal sensitivity.

Only on average 120 and 810 sub-samples from the full vector of 5,000 draws were necessary to achieve *r*^2^ *>* 0.99 between *C*-values for Joint analyses in three- and nine-spined sticklebacks, respectively (Fig. S7). This indicates that for empirical data *>* 1000 draws is not necessary but also that the results from three-spine sticklebacks are likely less sensitive to parameter choice compared to nine-spined sticklebacks, possibly due to the much stronger background levels of genetic differentiation in the latter.

In three-spined sticklebacks, detected outlier regions included three well-known inversions associated with marine–freshwater divergence (chromosomes 1, 11, and 21), as well as multiple regions on chromosome 4 encompassing the *EDA* locus (Colosimo et al., 2005). In addition, chromosomes 1, 4, 7, 12, and 20 contained four or more outlier regions, whereas chromosomes 13, 15, and 16 contained none (mean = 2.5 ORs per chromosome). In nine-spined sticklebacks, only chromosomes 7, 11, 20, and 21 contained more than one OR (maximum = 3), with six chromosomes containing no ORs (mean = 1.4 ORs per chromosome). These patterns are consistent with Fang et al. (2021), who reported 2.9 times more ORs in three-compared to nine-spined sticklebacks, although our analyses detected 1.9-fold and 2.2-fold higher numbers of ORs, respectively. Overall, the empirical results mirror simulation-based

## 4 Discussion

### 4.1 Overview and main contributions

In this study, we introduce and validate a genome-scan framework that explicitly integrates local linkage disequilibrium (LD), parameter uncertainty, and complementary inference methods to improve the detection of genomic regions under selection. Using forward-in-time simulations with known ground truth, we show that LD-scaling of association statistics substantially improves the recovery of true outlier regions across a wide range of demographic and selective scenarios, particularly under high gene flow where SNP-level approaches often fail. By aggregating evidence across parameter values and methods using a consistency-based score (*C*), the framework further (i) reduces sensitivity to arbitrary thresholds, (ii) suppresses false positives driven by background LD and (iii) prioritizes signals that persist under many plausible analytical choices. Re-analyses of empirical stickleback data demonstrate that these improvements translate to real genomes, recovering well-established regions of parallel adaptation with much greater coherence and reduced noise. Together, these results establish LD-aware, region-based, and uncertainty-integrated inference as a robust extension to existing genome-scan ap-proaches.

### 4.2 Why LD-aware, region-based inference improves genome scans

A central limitation of many genome-scan methods is the implicit assumption that loci represent independent tests, despite the pervasive influence of LD generated by demography, recombination heterogeneity, and selection itself (Felsenstein and Lewontin, 1975; Slatkin, 2008; Li et al., 2018). This mismatch contributes both to inflated false positives under complex population structure and to reduced power when stringent multiple-testing corrections are applied (Lotterhos and Whitlock, 2015; Hoban et al., 2016; Ravinet et al., 2017). Our results demonstrate that explicitly incorporating local LD structure into association statistics and evaluating inference at the level of genomic regions rather than individual SNPs substantially improves robustness across evolutionary scenarios.

Region-based inference is particularly well suited to forms of adaptation that generate extended LD footprints, including polygenic selection acting on clusters of loci, reduced effective recombination, and structural variants such as inversions (Kirkpatrick and Barton, 2006; Feder et al., 2012; Wellenreuther et al., 2019). In such contexts, elevated test statistics arise not only at causal variants but also at linked neutral loci, making SNP-level interpretation inherently noisy. Aggregating signal across linked markers therefore provides a biologically meaningful representation of selection that better reflects how adaptive processes manifest at the genomic scale.

### 4.3 Parameter uncertainty is intrinsic, not incidental

Across methods and simulation scenarios, variation in performance could be traced to how threshold parameters modulate the balance between true and false positives. Parameters defining outlier regions—such as minimum region size (*l*_min_) and clustering thresholds relative to LD decay (*τ*_LD*,ρ*_, *τ*_d*,ρ*_)—primarily influenced precision by filtering satellite clusters and over-fragmentation. In contrast, parameters controlling statistical stringency—such as *α* and the consistency threshold *τ*_C_—mainly re-duced false positives, often with comparatively smaller losses in recall. The LD window size (*w*) defined the spatial scale over which LD footprints were emphasized; too low values reduces the contrast between neutral and selected regions (increasing false positives) while too high values limit statistical power to only the highest local LD-signatures, for instance caused by inversions.

Crucially, no single parameter or threshold emerged as universally optimal. Instead, high performance consistently arose from combinations of parameter values, explaining the strong co-dependencies observed among thresholds. Partial *η*^2^ analyses further showed that while demographic parameters strongly shaped biological signals such as *Q_ST_ /F_ST_* and background LD, method choice primarily governed how those signals were partitioned into outlier regions. This distinction highlights that uncertainty in parameter choice is an inherent feature of genome scans rather than a technical inconvenience that can be eliminated by optimization.

### 4.4 Consistency-based integration and advances over LDna

The framework builds directly on earlier LD-based approaches such as LDna (Fang et al., 2021), sharing the core idea that selection should be inferred at the level of genomic regions rather than individual loci. The principal advance here is the replacement of genome-wide LD network construction with explicit LD-scaling of association statistics, combined with a formal consistency-based framework for integrating support across parameter draws and methods.

This study provides the first systematic validation of such an LD-based integration strategy using simulations with known ground truth. This allowed direct quantification of how LD-scaling, dimensionality reduction, and joint inference affect the balance between true and false positives across diverse demographic regimes. The stickleback re-analyses therefore serve primarily to illustrate the practical consequences of these improvements, demonstrating that LD-scaled and consistency-based joint analyses recover known targets of selection with substantially reduced noise while remaining conceptually aligned with LDna-based interpretations.

An additional practical advantage is computational efficiency. By restricting analyses to SNPs that already exceed association thresholds, the framework avoids constructing genome-wide LD networks among all loci, reducing both memory usage and runtime by several orders of magnitude compared to previous LDna-based workflows (Fang et al., 2020, 2021). Even in nine-spined sticklebacks—a system characterized by strong background differentiation—fewer than 1,000 parameter draws were sufficient to obtain highly reproducible *C*-scores, making the approach feasible for large genomic datasets to be analysed within a couple of hours.

### 4.5 Methodological interpretation and validation

The LD-scaled *F* ^′^ statistic relies on a quantile transformation that does not correspond to a previously described parametric test. In the context of genome scans, however, methodological validity is determined less by conformity to an analytical null distribution than by performance under realistic evolutionary conditions, where assumptions of marker independence are routinely violated by LD, demography, and recombination heterogeneity (e.g., Lotterhos and Whitlock, 2015; Hoban et al., 2016; Ravinet et al., 2017). The underlying intuition is that loci influenced by selection tend to be embedded within genomic regions showing elevated and spatially coherent LD, whereas background LD varies more gradually along chromosomes (Nielsen et al., 2005; Gompert et al., 2022). Weighting association statistics by local LD therefore amplifies signals consistent with selection-driven LD footprints, but also introduces systematic distortion of the null distribution because LD itself is spatially autocorrelated (Clifford et al., 1989). The quantile transformation explicitly addresses this by estimating and removing LD-induced inflation via permutation, while preserving excess signal that exceeds expectations under *H*_0_. Importantly, this procedure functions as a calibration step rather than a new hypothesis test, re-taining correct SNP ranking in the upper tail. Across simulations with known ground truth, LD-scaling consistently improved discrimination between true and false outlier regions, increased robustness to parameter choice, and enhanced recovery of spatially coherent LD footprints around causal loci. This performance-based validation provides a biologically meaningful justification for the transformation, independent of its analytical form.

An additional and important aspect of the quantile transformation is that the permutation procedure explicitly accounts for spatial autocorrelation in LD along chromosomes. By using circular permuta-tion of the *ld_w_* vector (Cabrera et al., 2012), the local correlation structure of LD is preserved while its alignment with the association statistic is randomized. This is critical because standard permuta-tion tests that ignore autocorrelation can substantially underestimate variance and inflate significance when applied to spatially structured genomic data (Clifford et al., 1989). At the same time, the present approach differs from classical permutation testing in an important practical respect: because the per-mutation is used solely to estimate a systematic distortion of the null distribution rather than to gen-erate empirical *p*-values, high reproducibility is achieved with very few permutations. In practice, ten permutations of *ld_w_* yielded nearly identical transformed statistics (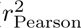 ≈ 1) across all simulated and empirical data sets. This contrasts sharply with conventional permutation frameworks, which of-ten require hundreds or thousands of replicates to achieve comparable precision. The combination of autocorrelation-aware permutation and rapid convergence substantially reduces computational cost, making LD-scaled calibration feasible for large genomic datasets. More generally, this strategy illus-trates how permutation can be repurposed as an efficient calibration tool rather than a brute-force sig-nificance test, with potential relevance for other analyses involving spatially or temporally autocorre-lated predictors beyond evolutionary genomics.

### 4.6 Simulation limits and localization accuracy

Simulated chromosomes were approximately one tenth the length of typical empirical maps, which in-flated long-range LD and increased the probability of hitchhiking at neutral loci. This design was nec-essary to detect LD signatures from relatively few segregating loci in the simulations, but it likely also contributed to the elevated false positive rates on QTN-bearing chromosomes. Although using LD-scaled distances to classify true versus false positives should partially mitigate this effect, all SNPs on these chromosomes were located within ∼10 cM of at least one QTN. In contrast, both because of the higher recombination rates and generally much larger effective population sizes, empirical genomes typically exhibit rapid LD decay over short physical distances providing higher mapping resolution(Gillespie, 2010; Hartl and Clark, 2007) . Our simulations are therefore conservative with respect to localization accuracy, suggesting that the separation between selected and neutral regions should be clearer in empirical datasets.

### 4.7 Empirical interpretation and conservative thresholding

For empirical analyses, we adopted *τ*_C_ = 0.2, close to the upper quartile of values observed among top-performing simulation draws, deliberately prioritizing robustness over maximal sensitivity. In combi-nation with stringent clustering thresholds, this choice reduced fragmentation of broad LD peaks and limited inflation of outlier-region counts. Several chromosomes contained no outlier regions at these thresholds. If these chromosomes lack QTNs, their absence of signal supports the interpretation that the thresholds are conservative yet appropriate. Simulations further showed that most false positives occurred on chromosomes already containing QTNs, indicating that chromosome-level patterns of out-lier regions are themselves informative even when precise causal variants cannot be resolved.

### 4.8 Scope, limitations, and practical guidance

Because *C* depends on the explored parameter space, absolute values should be interpreted relative to the sampled ranges. Narrower parameter ranges inflate *C*, whereas broader ranges emphasize robust-ness. Chromosome-level patterns provide an additional diagnostic: thresholds can be calibrated conser-vatively to avoid generating outlier regions on chromosomes that otherwise appear neutral.

The framework assumes that selection generates extended LD footprints and is therefore not intended as a replacement for GWAS or fine-mapping of isolated causal variants. Its strength lies in identifying coherent genomic regions shaped by selection, particularly in comparative contexts where robustness and interpretability are important. Outlier regions identified using Joint_C_ should therefore be viewed as hypotheses requiring independent validation.

### 4.9 Toward a new benchmark for outlier detection

Simulation-based benchmarking has long been recognized as the gold standard for evaluating genome-scan methods (Lotterhos and Whitlock, 2015; Lotterhos et al., 2022; Lotterhos, 2019). By extending this framework to region-level inference and integrating parameter uncertainty through AUC-PR^∗^, the present approach captures the spatial dependence among loci imposed by LD and reflects how selec-tion manifests at the genomic scale. We therefore propose that OR-level precision–recall metrics com-bined with uncertainty-aware evaluation should serve as a new benchmark for assessing outlier detec-tion methods in evolutionary genomics.

## Acknowledgements

We thank all members of the Guillaume lab who provided support and feedback during the develop-ment of this project, as well as X and Y for their comments on our manuscript. P.K. was supported by grant no. 354861 from the Research Council of Finland to F.G.

## Author contributions

PK developed the method and analysed the data with input from FG. PK and FG jointly planned, ex-ecuted, and interpreted the simulation experiments and wrote the manuscript.

## Data and code availability

All analyses were performed in R. The methods introduced here are implemented in an open-source R package, LDscanR, available at https://github.com/genomeinsights/LDscnR. Scripts to reproduce all simulations and empirical analyses will be submitted to https://figshare.com.

## Supplementary Information

**Figure S1:**
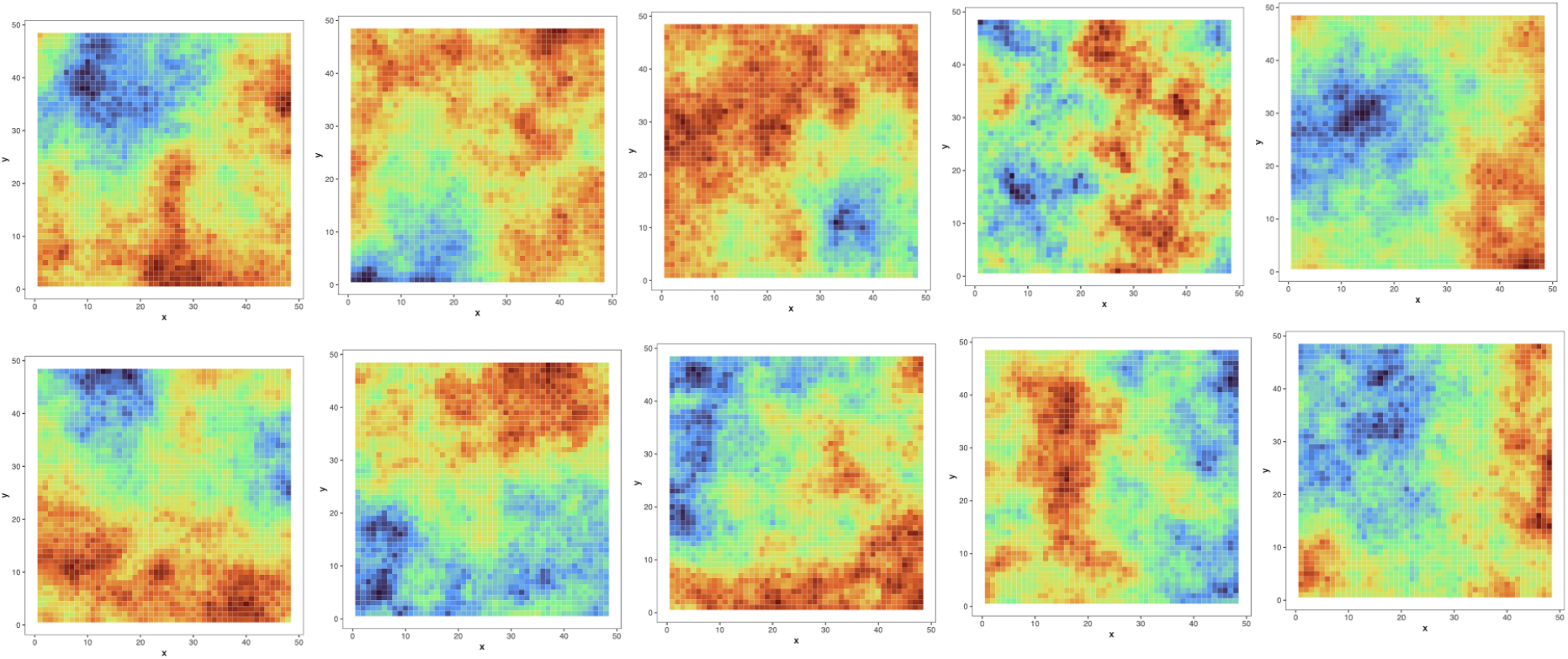
Simulated spatially autocorrelated environmental gradients used in forward simulations of local adaptation. Each panel represents one random realization of a continuous landscape on a 48 × 48 grid of demes, generated using an exponential variogram model (sill = 10, range = 50, nugget = 0.01). Colors indicate the rescaled environmental variable (range [0, 1]), which was subsequently used as input in Nemo. Replicate landscapes provide different random realizations of the same underlying spatial model, ensuring both realism and stochastic variation in environmental heterogeneity.

**Figure S2:**
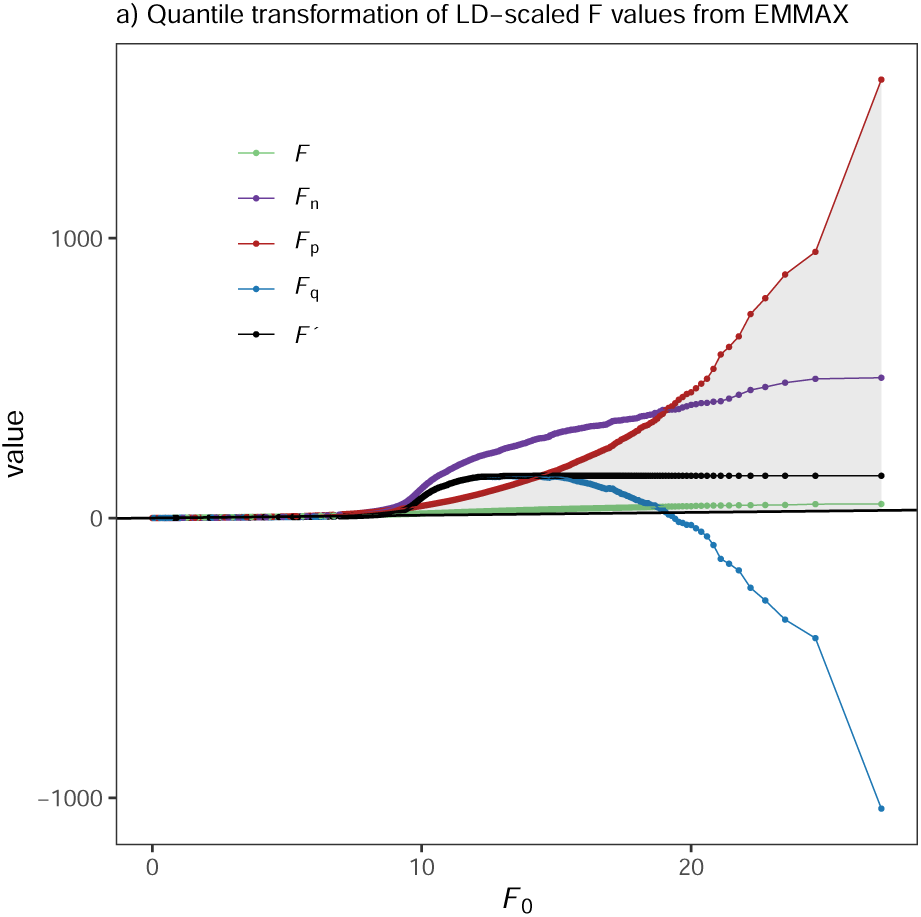
Quantile transformation of LD-scaled *F* -values in empirical three-spined stickleback data. *F* is the original test statistic from EMMAX. *F*_n_ = *ld_w_*× *F/̄F* represents the observed LD-weighted statistic. *F*_p_ is obtained by combining *F* with permuted *ld_w_* values to generate the null distribution. *F*_q_ is the quantile-transformed statistic obtained by adjusting *F*_n_ based on the difference between *Q*(*F*_p_) and the theoretical *F* -distribution (*F*_0_; shaded area). The final test statistic *F* ^′^ = cummax(*F*_q_) ensures proper SNP ranking in the upper tail.

**Figure S3:**
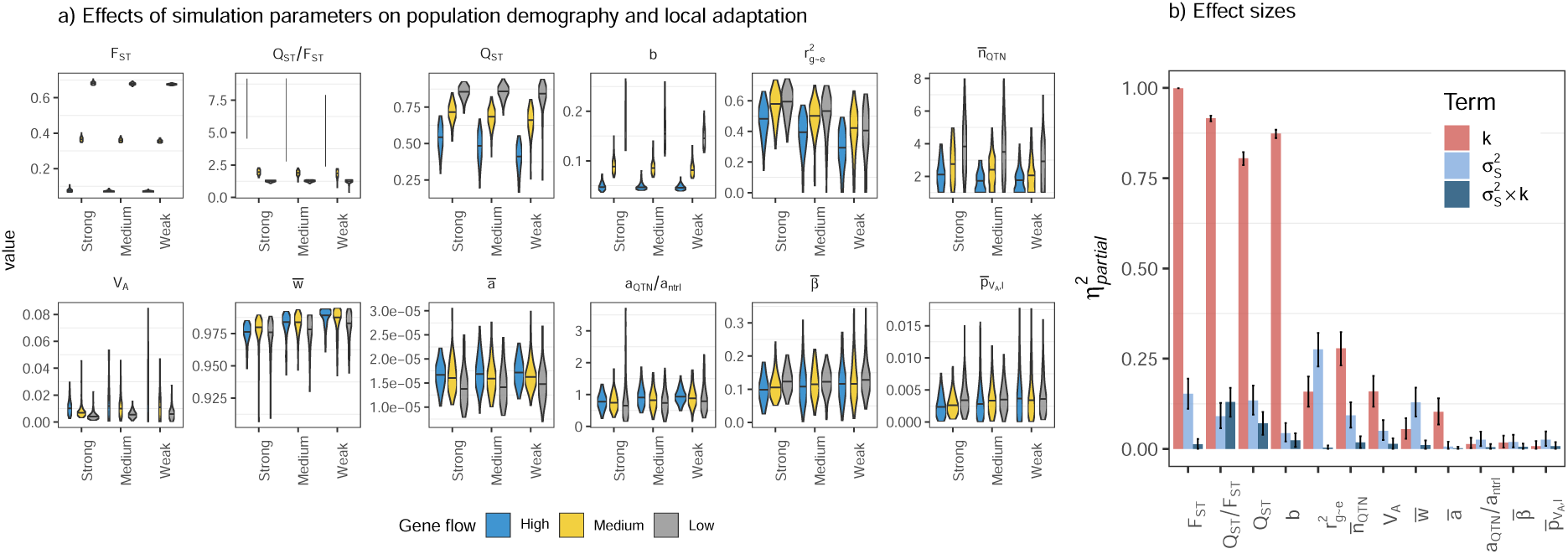
Effects of gene flow and selection intensity on population-level summary statistics in the simulations. (a) Violin plots showing the distribution of key demographic, genetic, and adaptive summary statistics across simulation replicates, stratified by gene flow (*k*) and selection intensity (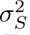). Me-dians are indicated by horizontal lines. Statistics include neutral and adaptive differentiation (*F_ST_* , *Q_ST_* , *Q_ST_ /F_ST_* ), additive genetic variance (*V_A_*), background linkage disequilibrium (*b*), LD-decay parameters (*a*), the number and effect size of segregating QTNs, and the strength of local adaptation 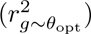. (b) Partial effect sizes (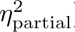) from Type-III ANOVA models quantifying the proportion of variance in each statistic explained uniquely by gene flow (*k*), selection intensity (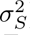), and their interaction. Bars represent point estimates and error bars denote 95% confidence intervals. Statistics are ordered by total explained variance to highlight which aspects of genomic structure and adaptation are most strongly shaped by simulation parameters.

**Figure S4:**
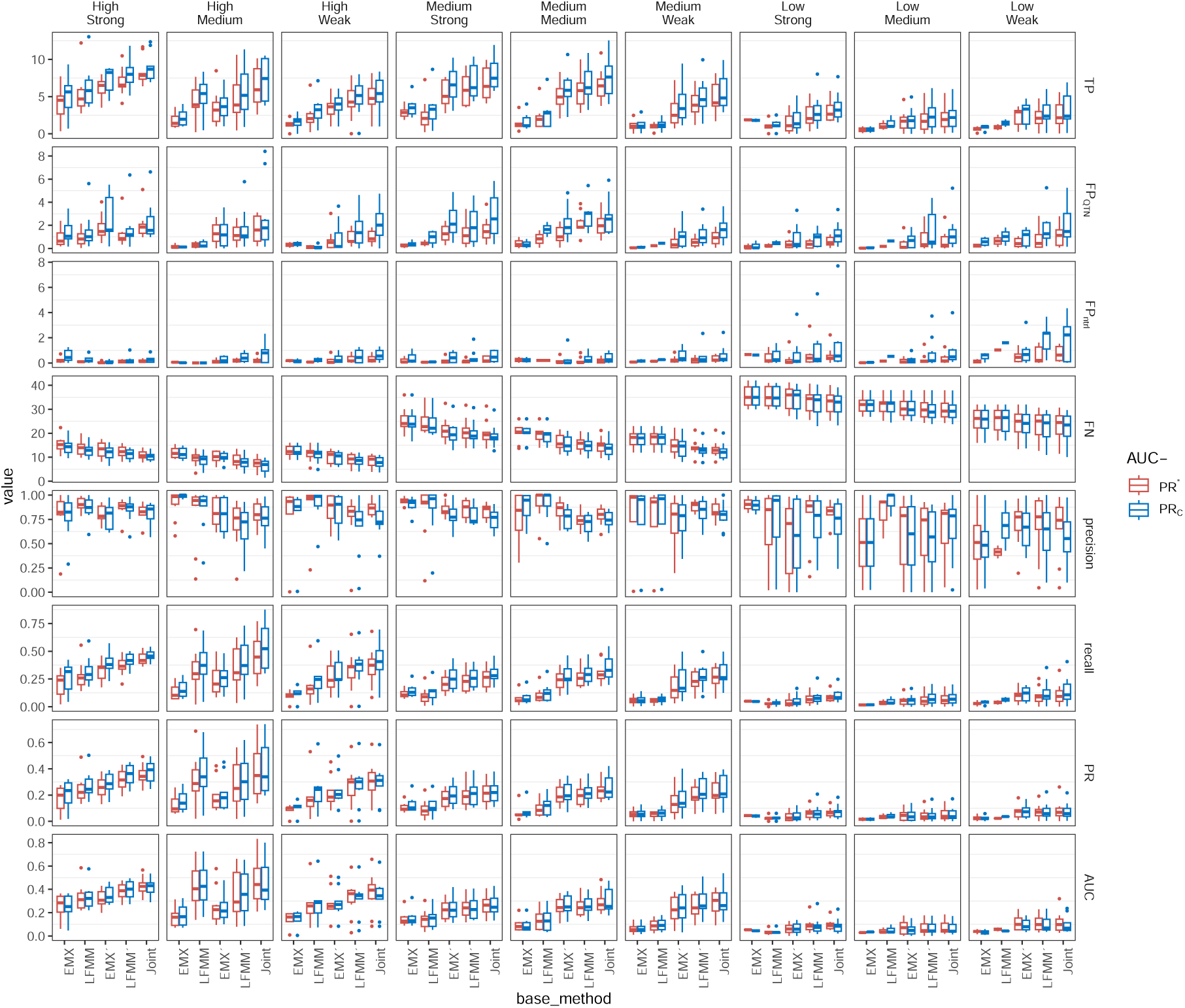
Summary of performance metrics from the top 10% (500/5,000) draws. Shows the distribution of the no. true positive ORs (TP), the no. false positive ORs separately for *QTN* (FP_QTN_) and *ntrl* (FP_ntrl_) chromosomes, the no. false negatives (FN), precision = TP*/*(TP + FP), recall = TP*/*(TP + FN), precision × recall (PR) and area under the PR curve (AUC). All values are means across simulated genomes (20 chromosomes simulated in the same selection landscape; *n* = 10), except AUC, which integrates performance across all draws. PR^∗^ and PR_C_ indicate whether draws are based (i) on all parameters (*α*, *l_min_*, *w*, *τ_LD_* and *τ_d_*) or, (ii) on the composite *C*-score that summerises the information from *α*, *l_min_* and *w* from (i). While PR indicates overall higher performance for PR_C_ than PR^∗^ (second panel from the bottom), when integrating across all draws (not just the top 10%), their performances are indistinguishable from each other (AUC, bottom panel, see main document for details).

**Figure S5:**
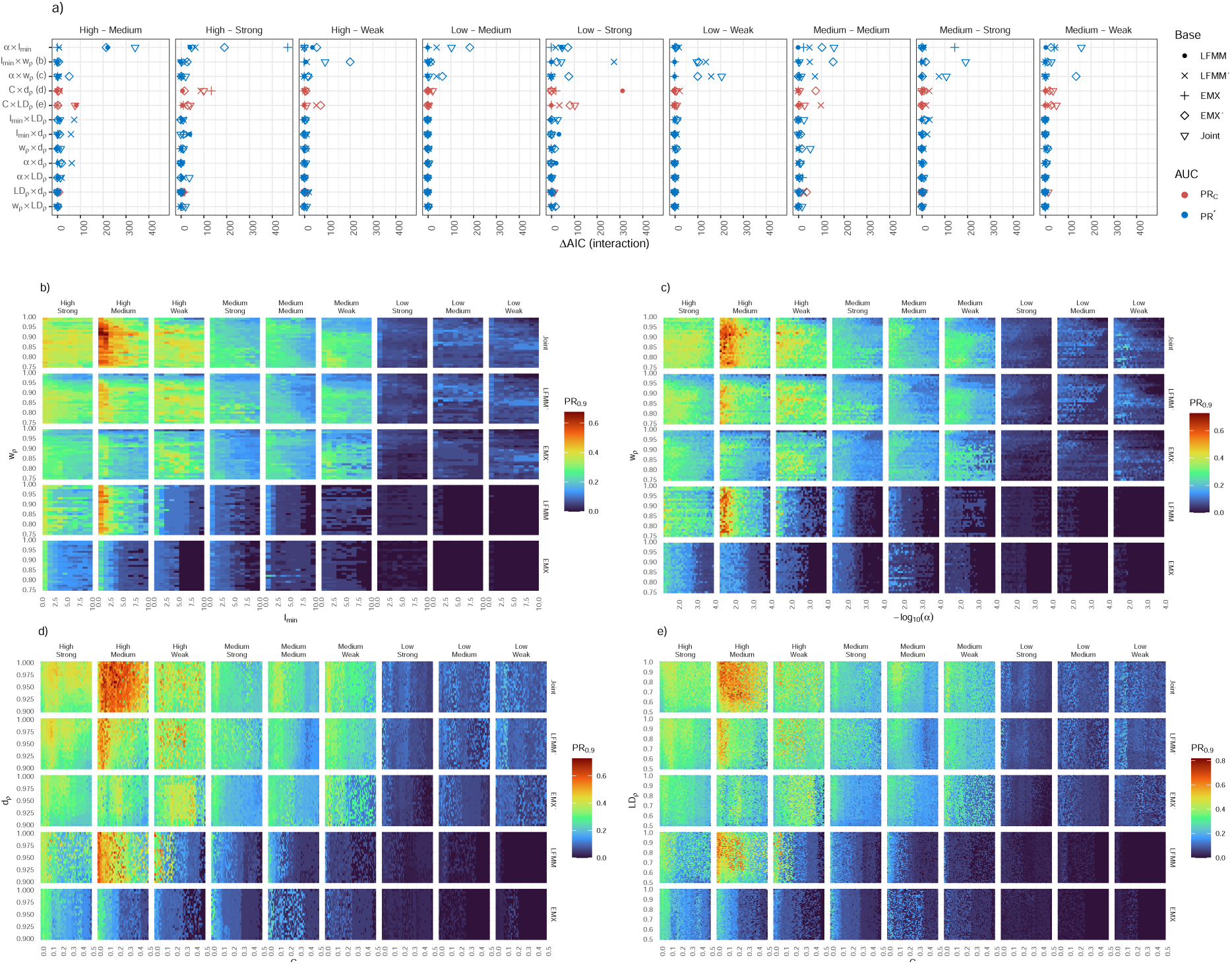
Heatmaps illustrating co-dependencies among threshold parameters across outlier methods and simulation scenarios. (a) Summary of evidence for pairwise parameter interactions, quantified as the improvement in model fit when including an interaction term (ΔAIC; additive model vs. interaction model) in logistic regressions predicting whether a parameter draw fell within the top 10% of PR (i.e., *I*_top_ = 1). Points are shown for each base method (shape) and AUC implementation (color), and parameter pairs are ordered from strongest to weakest average interaction support (top to bottom). (b–c) Heatmaps of PR_0.9_ (90th percentile of PR across draws) for the four parameter pairs with the strongest interaction support in panel (a), illustrating how high performance is achieved along ridges in joint parameter space rather than at a single optimum. The *α* × *l*_min_ heatmap is shown separately in Fig. 6b.

**Figure S6:**
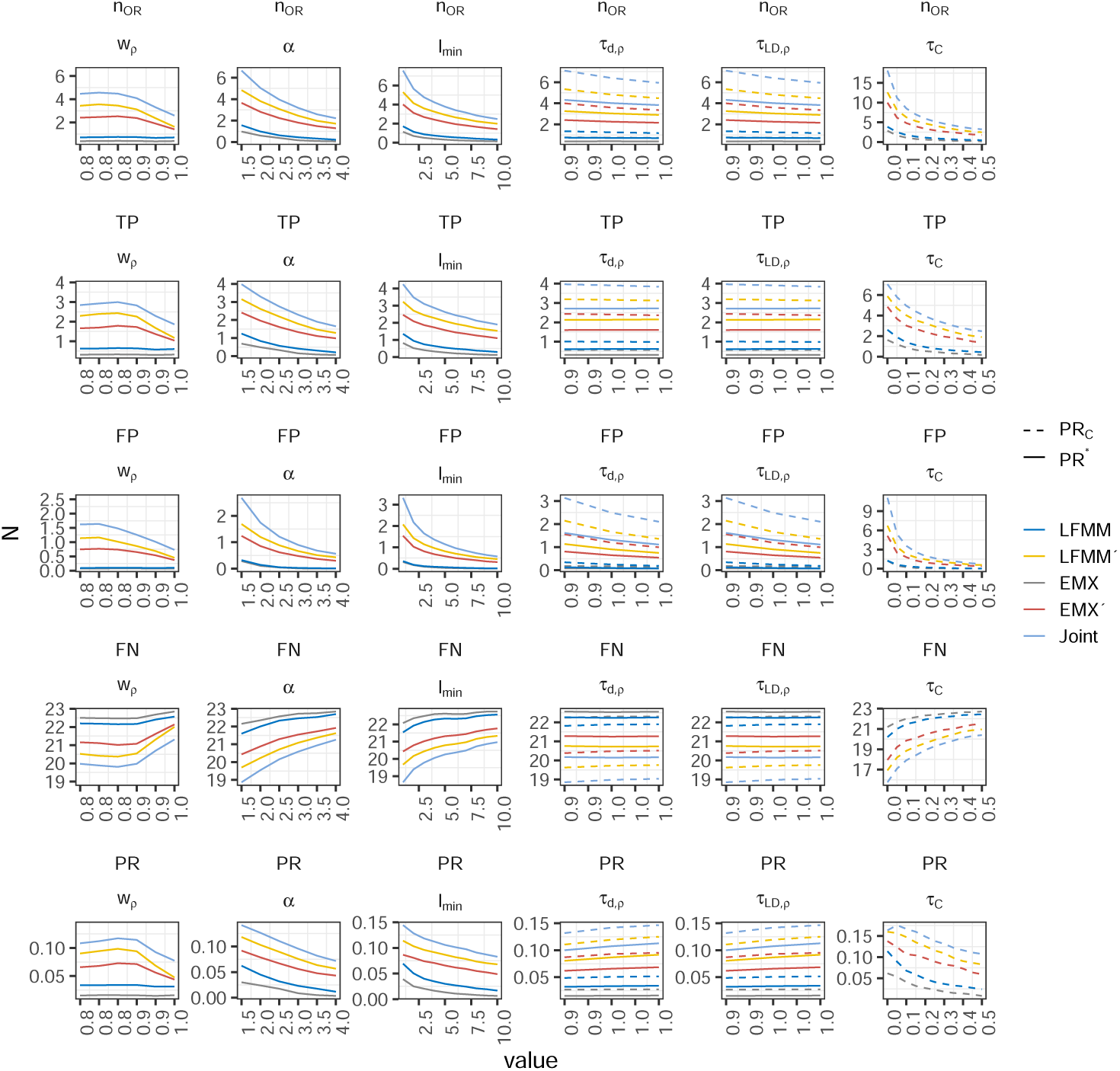
Marginal effects of threshold parameter choice across all 5,000 random draws on outlier detection outcomes. Panels summarize the distribution of the number of outlier regions (*n*_OR_), true positives (TP), false positives (FP), false negatives (FN), and the precision–recall product (PR) as a function of each threshold parameter. Thresholds are shown on their sampling scales, with *α* expressed as — log_10_(*α*). Across parameters, increasing threshold stringency generally reduces the number of false positives, often also at the cost of reduced recall.

**Figure S7:**
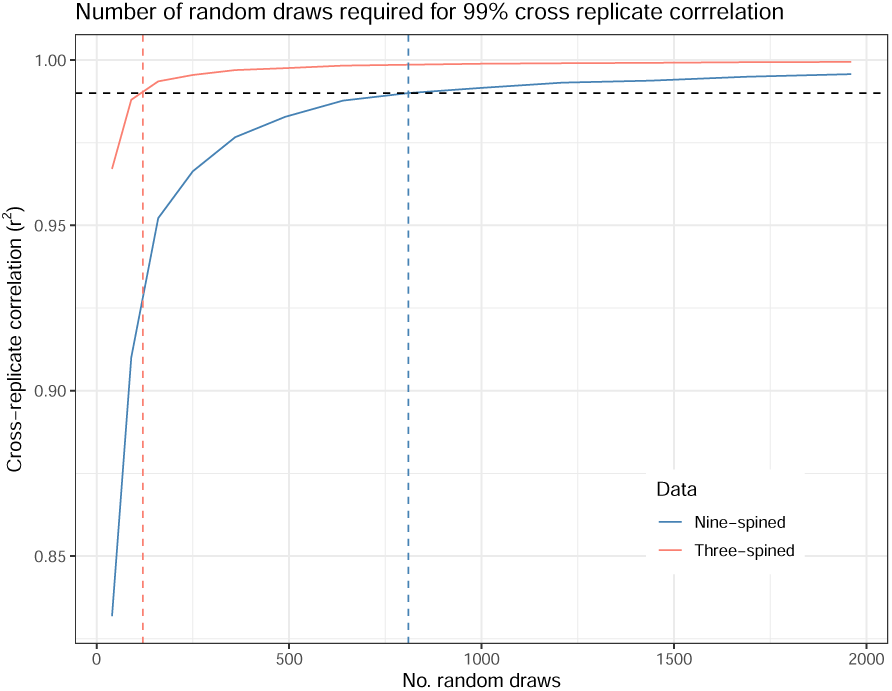
Convergence of *C*-scores with respect to the number of random parameter draws in empirical stickleback data. The figure shows the mean squared correlation coefficient (*r*^2^; *n* = 100 resamples) between *C*-scores estimated from a subset of parameter draws and those obtained from the full set of 5,000 draws, plotted as a function of the number of draws subsampled with replacement. Results are shown separately for three- and nine-spined sticklebacks. A threshold of high cross-replicate consistency (*r*^2^ *>* 0.99) was reached at approximately 120 and 810 draws for three- and nine-spined sticklebacks, respectively, indicating that substantially fewer draws than used in the simulations are sufficient to obtain stable *C*-score estimates in the empirical datasets.

**Table 1:** Summary of mathematical variables, parameters, and derived quantities used throughout the study.

| Symbol | Type | Description |
| --- | --- | --- |
| **Landscape, Demography, and Dispersal** |  |  |
| $K_d$ | parameter | Carrying capacity per deme (fixed). |
| $K_{\text{tot}}$ | parameter | Total carrying capacity of the landscape. |
| $m_{ij}$ | random variable | Probability of dispersal from deme $i$ to deme $j$. |
| $s_{ij}$ | parameter | Physical distance between demes. |
| $a_m$ | parameter | Mean dispersal scale parameter in the Gaussian kernel. |
| $\parallel$ | parameter | Shape parameter determining dispersal kernel heaviness. |
| **Recombination Map and Genomic Coordinates** |  |  |
| cM | unit | CentiMorgan (recombination distance). |
| bp | unit | Base-pair position. |
| **Genetic Effects and Trait Architecture** |  |  |
| $\beta_l$ | parameter | Additive allelic effect size at QTN $l$. |
| $g$ | random variable | Individual genotypic value (sum of allelic effects). |
| $\beta_{l,x}, \beta_{l,y}$ | random variable | Maternal and paternal allelic effects at locus $l$. |
| $N$ | parameter | Number of QTNs used in variance decomposition. |
| $p_l$ | random variable | frequency of ancestral allele at QTN $l$. |
| $p_{V_A,l}$ | derived quantity | Proportion of additive genetic variance explained by QTN $l$ on a chromosome. |
| **Environmental Optima and Fitness** |  |  |
| $\theta_{\text{opt}}$ | parameter | Local phenotypic optimum defined by the landscape. |
| $W$ | random variable | Individual fitness. |
| $\sigma_S^2$ | parameter | Selection variance (inverse of the strength of selection). |
| **Linkage Disequilibrium and Decay Model** |  |  |
| *Continued on next page* |  |  |

**Table 1 (continued)**
| Symbol | Type | Description |
| --- | --- | --- |
| $\rho$ | parameter | LD-decay quantile used for threshold normalization. |
| $r^2, r_\rho^2$ | derived quantity | Linkage disequilibrium between a pair of SNPs (bps, LD-quantile). |
| $d, d_\rho$ | derived quantity | Physical distance between SNPs (bps, LD-quantile). |
| $a$ | parameter | LD decay rate (nonlinear model). |
| $b$ | parameter | Background LD level. |
| $c$ | parameter | LD at distance zero (95% quantile). |
| **Association Testing and Test Statistics** |  |  |
| $F$ | random variable | Association statistic from LFMM or EMMAX. |
| $F_n$ | derived quantity | LD-weighted and normalized $F$. |
| $F_p$ | derived quantity | Permutation-based null statistic using permuted $ld_w$. |
| $F_q$ | derived quantity | Quantile-corrected statistic accounting for heavy tails. |
| $F'$ | derived quantity | Final LD-scaled test statistic: $\text{cummax}(F_q)$. |
| $ld_w$ | derived quantity | Median $r^2$ within LD window of size $w$. |
| $w, w_\rho$ | parameter | LD window size (bps, LD-quantile). |
| **Outlier Region (OR) Detection** |  |  |
| $\tau_{LD}, \tau_{LD,\rho}$ | parameter | LD threshold for clustering SNPs into ORs (bps, LD-quantile). |
| $\tau_d, \tau_{d,\rho}$ | parameter | Maximum physical distance allowed within an OR (bps, LD-quantile). |
| $l_{\min}$ | parameter | Minimum SNP count to define an OR. |
| $QTN_f$ | derived quantity | Focal QTN assigned to an OR. |
| $\alpha$ | parameter | Significance threshold for outlier identification. |
| $C$ | derived quantity | Consistency score for SNP-level recurrence across parameter draws. |
| $\tau_C$ | parameter | Threshold on consistency score for OR detection. |

Table 1 (continued)
| Symbol | Type | Description |
| --- | --- | --- |
| **Evaluation Metrics** |  |  |
| TP, FP, FN | derived quantities | True, false, and missed positives at OR level. |
| Precision | derived quantity | $TP/(TP + FP)$. |
| Recall | derived quantity | $TP/(TP + FN)$. |
| PR | derived quantity | Precision × Recall. |
| AUC-PR* | metric | Area under PR* across parameter sets. |
| AUC-PR<sub>C</sub> | metric | Area under PR<sub>C</sub> with dimensionality reduction. |
| $\mathcal{R}$ | metric | Robustness score: AUC-PR/$\max(\text{PR})$. |

